# Genetic code expansion in the engineered organism Vmax X2: High yield and exceptional fidelity

**DOI:** 10.1101/2021.06.22.449487

**Authors:** Sebasthian Santiago, Omer Ad, Bhavana Shah, Zhongqi Zhang, Xizi Zhang, Abhishek Chatterjee, Alanna Schepartz

## Abstract

We report that the recently introduced commercial strain of *V. natriegens* (Vmax X2) supports robust unnatural amino acid mutagenesis, generating exceptional yields of soluble protein containing up to 5 non-canonical α-amino acids (ncAA). The isolated yields of ncAA-containing superfolder green fluorescent protein (sfGFP) expressed in Vmax X2 are up to 25-fold higher than those achieved using commercial expression strains (Top10 and BL21) and more than10-fold higher than those achieved using two different genomically recoded *E. coli* strains that lack endogenous UAG stop codons and release factor 1 and have been optimized for improved fitness and preferred growth temperature (C321.ΔA.opt and C321.ΔA.exp). In addition to higher yields of soluble protein, Vmax X2 cells also generate proteins with significantly lower levels of mis-incorporated natural α-amino acids at the UAG-programmed position, especially in cases where the ncAA is an imperfect substrate for the chosen orthogonal aminoacyl tRNA synthetase (aaRS). This increase in fidelity implies that use of Vmax X2 cells as the expression host can obviate the need for time-consuming directed evolution experiments to improve specific activity of highly desirable but imperfect ncAA substrates.

## Introduction

Non-canonical amino acid mutagenesis, often referred to as genetic code expansion (GCE), is a powerful tool for introducing unique chemical functionality or reactivity into an expressed protein.^1–3^ Literally hundreds of diverse non-canonical α-amino acids (ncAA), including those that support protein labeling or conjugation, can be introduced into proteins^3–6^ biosynthesized in laboratory *E. coli* strains, pathogenic^7^ and soil bacteria,^8^ yeast,^9^ mammalian cells,^9–11^ and whole organisms.^12–14^ Unnatural amino acid mutagenesis *in vivo* demands co-expression of one or more aminoacyl-tRNA synthetase (aaRS)/tRNA pairs that are orthogonal in the expression host. Commonly used pairs include variants of pyrrolysyl aminoacyl-tRNA synthetase (PylRS)/pylT from Methanosarcina,^15–19^ tyrosyl aminoacyl-tRNA synthetase (TyrRS)/tRNA^Tyr^ from *Methanococcus jannaschii*^20–23^ and others;^24–26^ newly identified orthogonal pairs include those from *Lumatobacter nonamiensi, Sorangium cellulosum*, and *Archaeoglobus fulgidus*.^24^

In addition to its utility in basic research, non-canonical amino acid mutagenesis has significant and growing importance in the pharmaceutical and biotechnological industries. The ability to efficiently introduce reactive bioorthogonal functionality into a therapeutic antibody provides a streamlined route to homogeneous antibody-drug conjugates with high (∼95%) conjugation efficiency.^27–29^ Incorporation of non-canonical α-amino acids can be leveraged to confer favorable therapeutic properties, such as increased circulation half-life and improved bioactivity.^30–32^ Finally, GCE expands the chemical space that is accessible for the development of novel macrocycles and therapeutic peptides.^33–35^

For almost all genetic code expansion applications, yield and purity are paramount.^36^ One factor that can limit the yield of a protein carrying one or more ncAAs is competition between the mis-acylated suppressor tRNA and release factor 1 (RF1), both of which recognizes the amber UAG stop codon. Recognition of the mis-acylated suppressor tRNA leads to incorporation of the ncAA, whereas recognition by RF1 triggers early translation termination.^37,38^ Another factor that can limit yields is that the orthogonality of an aaRS/tRNA pair is rarely absolute, resulting in suppressor tRNAs that are acylated incorrectly with one or more α-amino acids. To circumvent this issue, researchers have developed *E. coli* strains that lack RF1, including those that have been genomically recoded to eliminate all or some of the 321 endogenous UAG stop codons in *E. coli*.^39–41^ While these strains can improve expression yields, genomically recoded organisms that lack RF1 suffer from fitness defects as well as higher levels of misincorporation events when utilizing suboptimal aaRS/tRNA pairs.^42–45^ Cell-free translation systems offer the opportunity to omit RF1 as well as tune the individual levels of near-cognate tRNAs to potentially increase incorporation fidelity, but these systems are significantly more costly than cellular bioproduction.^46,47^

Further challenges arise when high-yield expression of the target protein in *E. coli* demands low temperatures. Many orthogonal synthetases are derived from thermophilic organisms and exhibit minimal activity at temperatures below 25°C.^48,49^ The requirement for higher expression temperatures can lower the yield of target proteins that are unstable and/or insoluble under these conditions. *Vibrio natriegens*, a bacteria isolated originally from salt marsh mud, is the fastest growing organism on record and expresses many recombinant proteins in exceptional yields and at a variety of temperatures (20°C – 37°C).^50–56^ Previous research has revealed considerable compatibility between *E. coli* and *V. natriegens* in terms of commonly used genetic elements (i.e. promoters, ribosome binding sites, etc.) and plasmids.^52,57^ This compatibility facilitates the use of numerous extensively optimized *E. coli* GCE plasmid systems with little or no plasmid modification. Vmax X2 cells also possess advantages for the production of protein destined for used in animals, where endotoxin contamination remains a persistent concern, generating endotoxin titers even lower than those seen in cells like ClearColi^®^ cells.^58^

Here we demonstrate that the recently introduced commercial strain of *V. natriegens* (Vmax X2) supports robust unnatural amino acid mutagenesis, generating exceptional yields of soluble protein containing up to 5 ncAAs.^59^ Yields are especially high when ncAA are introduced using the *M. jannaschii* tyrosyl aminoacyl-tRNA synthetase (TyrRS)/tRNA^Tyr^ variant pCNFRS.^20,21^ The isolated yields of ncAA-containing superfolder green fluorescent protein (sfGFP) expressed in Vmax X2 are up to 25-fold higher than those achieved using commercial expression strains (Top10 and BL21) and more than 10-fold higher than those achieved using two different genomically recoded *E. coli* strains that lack endogenous UAG stop codons and have been optimized for improved fitness and preferred growth temperature (C321.ΔA.opt and C321.ΔA.exp, Addgene strains #87359 and #49018).^39,40^ The rapid doubling time of Vmax X2 (∼10-14 min)^52^ also translates into a highly convenient three-day workflow for protein expression, as opposed to the traditional 4-day workflow for protein expression using traditional *E. coli* strains. In addition to high yields, Vmax X2 cells also generate proteins with significantly lower levels of mis-incorporated natural α-amino acids at the UAG-programmed position, especially in cases where the ncAA is an imperfect substrate for the chosen orthogonal aminoacyl tRNA synthetase (aaRS). Thus, use of Vmax X2 can obviate the need for time-consuming directed evolution experiments to improve the specific activity of highly desirable but imperfect aaRS substrates.

## Results

To evaluate Vmax X2 as a host organism for unnatural amino acid mutagenesis, we first asked if it would support the incorporation of a single ncAA into sfGFP using two orthogonal translation systems (OTS) that are used commonly in *E. coli*. The first is the *p*-cyano-L-phenylalanyl aminoacyl-tRNA synthetase (pCNFRS)-tRNA_CUA_^Tyr^ pair^20,21^ derived from *M. jannaschii*, while the second is the pyrrolysyl-tRNA synthetase (PylRS)-tRNA_CUA_^Pyl^ pair^15^ derived from *M. mazei*. Both of these aaRS/tRNA pairs support the incorporation of chemically diverse ncAA into proteins in *E. coli* but only the activity of the PylRS)-tRNA_CUA_^Pyl^ pair^15^ has been tested in *Vibrio natriegens*.^59^ Specifically, we asked whether either of these aaRS/tRNA pairs would support the incorporation of *p*-azido-L-phenylalanine (using pCNFRS) or Boc-L-Lysine (using PylRS) at position 2 of sfGFP (Figure 1A). Vmax X2 cells were transformed with either pEVOL-mmPyl or pEVOL-CNF^60^ (along with pET-S2TAG sfGFP^61^) and grown in Vmax-optimized media supplemented with either 0.5 mM 4-azido-L-phenylalanine (pAzF) or 10 mM Boc-L-lysine (BocK)^62^ at a temperature of 37°C. Vmax X2 cells grew quickly under these conditions, reaching saturation after approximately 4 hours (**Figure 2A** and **Supplementary Figure 1**).

**Figure 1.**
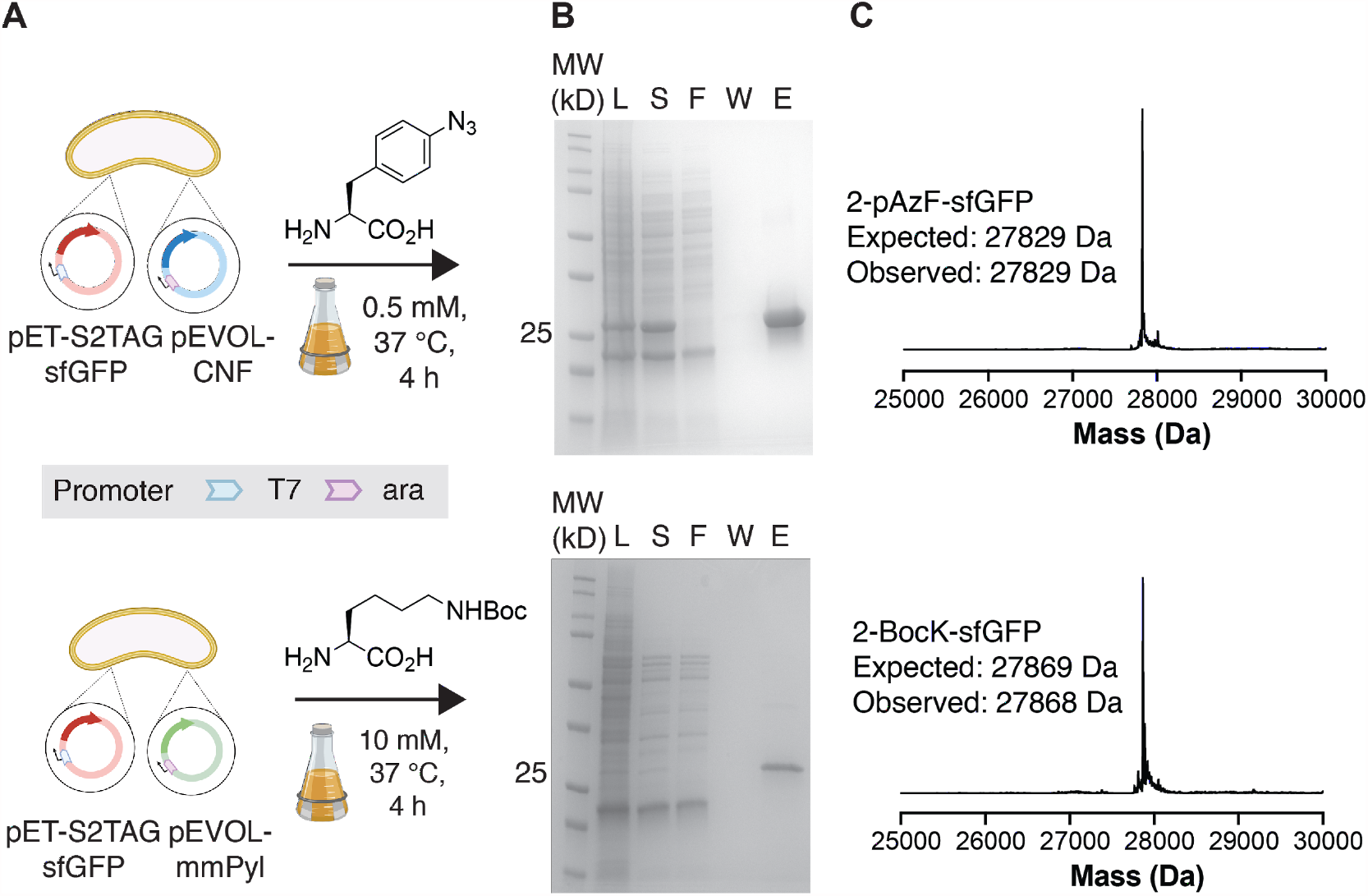
Genetic code expansion in Vmax X2. (A) Vmax X2 cells were transformed with pET-S2TAGsfGFP^61^ and either pEVOL-CNF^60^ or pEVOL-mmPyl to induce expression of sfGFP bearing a ncAA at the second position of sfGFP. Cells were induced and incubated for 4 hours at 37°C in the presence of 0.5 mM *p*-azido-L-phenylalanine (pAzF) (*p*CNFRS) or 10 mM L-Boc-lysine (BocK) (MmPylRS). (B) SDS-PAGE gels illustrate proteins produced in Vmax X2 cells transformed with pET-S2TAGsfGFP and either pEVOL-CNF or pEVOL-mmPyl. L = lysate; S = supernatant; F = flow-through; W = wash; E = elution. (C) Intact protein mass spectra of 2TAG sfGFP variants purified from Vmax X2 cells co-expressing *p*CNFRS (top) or MmPylRS (bottom) in the presence of pAzF or BocK, respectively.

**Figure 2.**
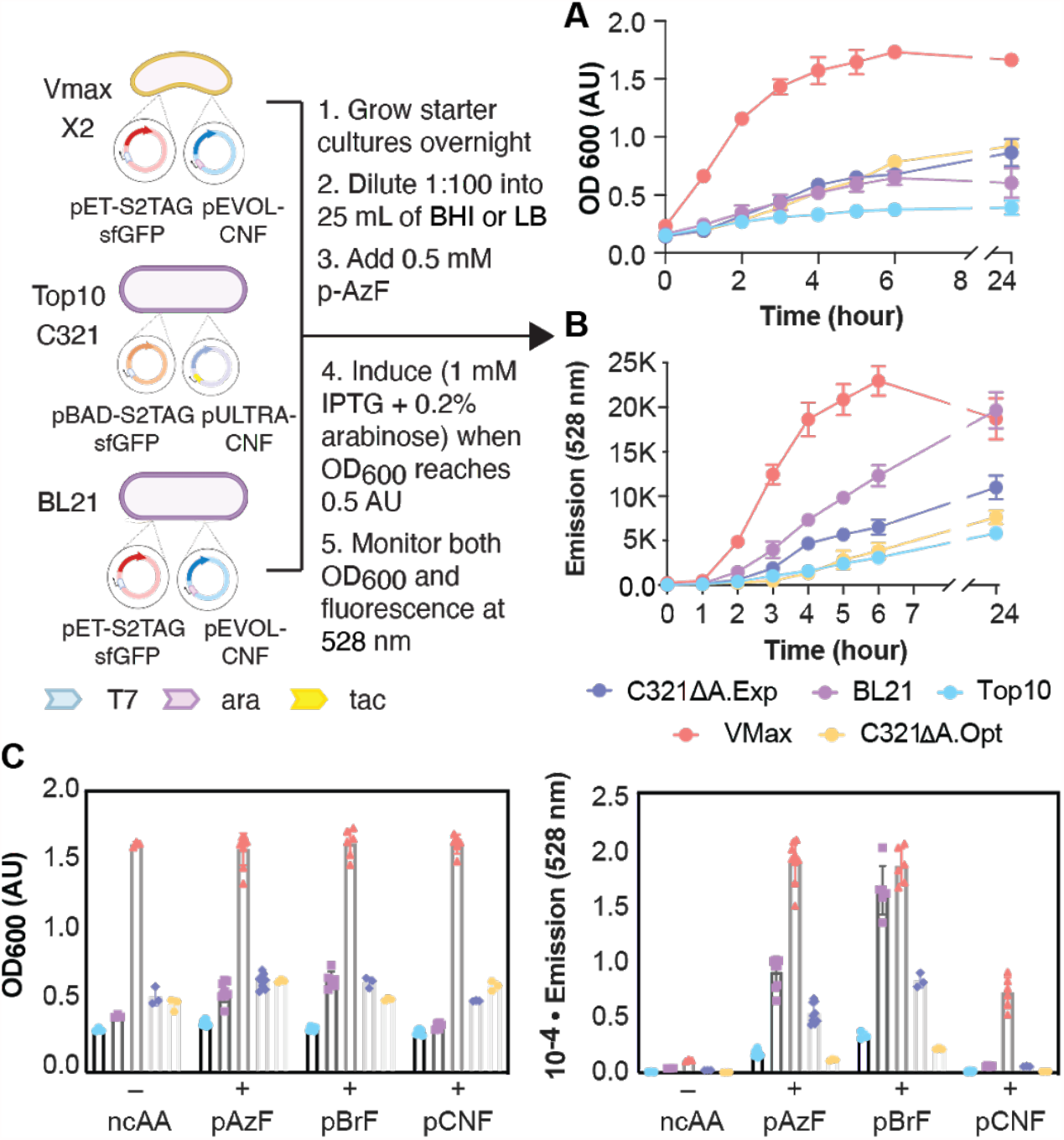
Growth and sfGFP expression in Vmax X2 versus traditional (Top10, BL21) and genomically recoded (C321)^39,40^ *E. coli* strains. (A) Plot of the OD_600_ of each cell growth as a function of time. Vmax X2 and BL21 cells were transformed with pET-S2TAGsfGFP and pEVOL-CNF, whereas Top10 and C321 cells were transformed with pBAD-S2TAGsfGFP and pULTRA-CNF (Top10, C321) to induce expression of sfGFP bearing a ncAA at the second position of sfGFP. After induction, cells were grown for 24 hours at 37°C (Vmax X2, BL21, Top10, C321.ΔA.exp) or 34°C (C321.ΔA.opt) in the presence of 0.5 mM pAzF. (B) Plot of the emission of each cell growth at 528 nm as a function of time. (C) Plots comparing the OD_600_ and 528 nm fluorescence of each growth at the 4 h time point in the presence or absence of pAzF, *p*-bromo-L-phenylalanine (pBrF), or *p*-cyano-L-phenylalanine (pCNF).

The 2TAG sfGFP variants expressed in Vmax X2 during a 4-hour incubation were isolated using immobilized metal affinity chromatography (IMAC) and their purity and identity assessed using SDS-PAGE and mass spectrometry (**Figure 1B and C**). In each case, SDS-PAGE evaluation of the isolated protein products revealed a prominent band just above the 25 kDa marker as expected, whose intact mass spectrum was consistent with incorporation of a single copy of either BocK or pAzF. To verify that the non-canonical α-amino acid was introduced into the expected (2^nd^) position, the proteins produced in Vmax X2 cells were digested with the endoproteinase Glu-C and analyzed further by LC-MS/MS (**Supplementary Figure 2**). Sequence matching of the digested peptides confirmed the incorporation of BocK and pAzF at position 2 of sfGFP. These data indicate that the orthogonal synthetases pCNFRS and PylRS are expressed and active in Vmax X2 cells, acylate their cognate tRNAs with the provided ncAA, and that the charged tRNA is utilized by the Vmax X2 translational machinery. The yield of sfGFP containing an ncAA at position 2 was 8.8 mg (BocK) and 387.2 mg (pAzF) per liter of culture. The yield of sfGFP containing BocK at position 2 is more than 8-fold higher than that observed previously (∼1 mg/L), likely due to the use of a stronger promoter for the sfGFP transcript, as well as ncAA-dependent effects.^59^

Next, we set out to evaluate how the yield of ncAA-containing sfGFP produced in Vmax X2 cells compared to those obtained in several *E. coli* strains used for protein expression and genetic code expansion. The strains evaluated included Top10 (a broad utility strain related to DH10B™), BL21 (optimized for protein expression from T7 promoters), as well as two genomically recoded strains, C321.ΔA.exp^39^ and C321.ΔA.opt,^40^ in which the 321 endogenous UAG stop codons are replaced by the alternative stop codon UAA. Both GROs are derived from the original C321.ΔA; the former has been engineered to lower the rate of spontaneous mutagenesis and can be grown at 37 °C; the latter carries additional mutations that improve doubling time. Because C321 strains lack T7 RNA polymerase,^39^ sfGFP expression in these strains is under control of the commonly used pBAD promoter. All strains were grown under their own optimized conditions and in the presence of 0.5 mM pAzF; both OD_600_ and fluorescence at 528 nm (λ_max_ for sfGFP) were monitored as a function of time (**Figure 2A**).

As expected, the Vmax X2 cultures grew faster than all others and reached saturation at an OD_600_ of 1.5 approximately 4 hours after induction. All other strains required more than 24 h to reach an OD_600_ of 1.0. As judged by the emission value at 528 nm, sfGFP expression in Vmax X2 reached a maximal value after 6 hours of expression and then decreased slightly. In the case of all other strains, the signal at 528 nm increased linearly over time over the entire course of the experiment (24 h). The greatest difference in OD_600_ and fluorescence occurred at the 4h time point (**Figure 2A**).

Examination of the OD_600_ and emission at 528 nm after 4 h incubation in the presence of three different pCNFRS substrates (pAzF, pBrF, and pCNF) reveals several trends (**Figure 2C**). First, as expected, the Vmax X2 growth rate exceeded that of any other strain in the absence of a ncAA or in the presence of 0.5 mM pAzF, pBrF, or pCNF. The presence or the identity of the ncAA had little or no effect on the growth rate of any strain examined. The changes in fluorescence at 528 nm show more significant changes and greater dependence on ncAA identity. For example, although the values for 528 nm emission in the presence of pAzF mirrored the OD_600_ values across all strains, the values in the presence of pBrF were higher than expected in BL21. The values for 528 nm emission in the presence of pCNF were low in all strains other than Vmax X2. Despite these differences, two overarching trends emerge: in all cases, the 528 nm emission values suggest that higher levels of sfGFP are produced in Vmax X2 than in any other strain tested, and that the yields in BL21 cells exceed those obtained in either C321.ΔA.exp^39^ or C321.ΔA.opt.^40^ A plot of 528 nm fluorescence/OD_600_ (**Supplementary Figure 1C**) shows that Vmax X2 cells often express less sfGFP per cell, except in the case where pCNF was incorporated, compared to BL21 and C321.ΔA.exp.

Although fluorescence emission at 528 nm is often taken as a measure of sfGFP expression, for most applications it is the isolated, purified protein yield that matters more, and isolated yields can be affected negatively if aggregation occurs at high protein concentration.^63^ To evaluate whether the isolated yields would parallel fluorescence at 528 nm, we isolated sfGFP from each strain after a 4 h growth (**Figure 3A** and **Supplementary Table 1**). The unoptimized yield of isolated sfGFP produced in Vmax X2 cells was at least 2-fold and in some cases as much as 25-fold higher than the yield obtained in any *E. coli* strain tested. In the case of pAzF, the yield of sfGFP produced in Vmax was 25-fold higher than the yield obtained in either Top10 or C321.ΔA.opt cells. In fact, although proteins yields can be affected negatively by high expression titers, for Vmax X2 the opposite trend is observed: significantly more soluble protein is obtained per unit 528 nm absorbance in Vmax X2 than in any other strain, across all ncAA examined (**Supplementary Figure 3**).

**Figure 3.**
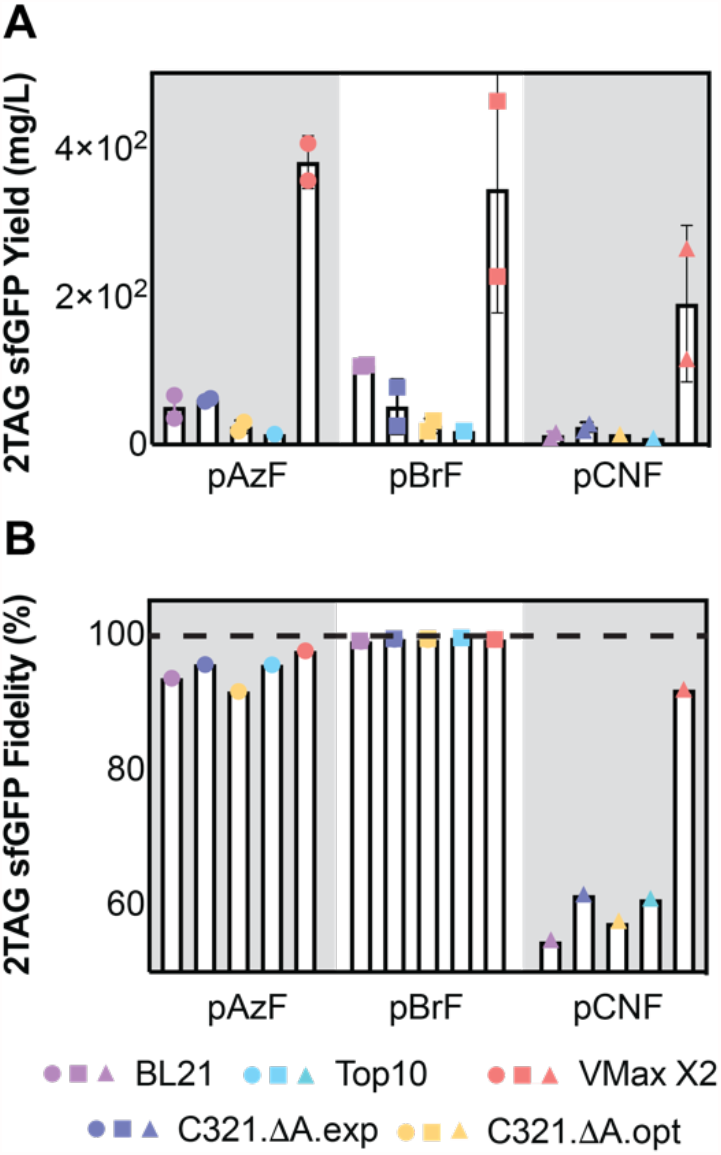
Yield and purity of sfGFP containing a single ncAA when expressed in Vmax X2 versus *E. coli* strains. (A) Isolated yield (mg/L) and (B) fidelity (%) of sfGFP containing the indicated ncAA at position 2 when expressed in the indicated strain. Cells were grown for 4 h, lysed *via* sonication, and sfGFP was isolated using IMAC. Yields were determined using the Pierce 660 nm Protein Assay (Thermo Scientific) and a BSA standard curve. Fidelity was determined from LC-MS/MS data as the fraction of the Glu-C-generated N-terminal peptide MXKGEE containing the desired ncAA (X) at position 2 relative to all other detectable amino acids at that position. The dashed line indices 100% fidelity. Additional LC-MS/MS data is found in **Supplementary Figures 4 and 5**.

One strain-dependent complication of GCE is misincorporation of an endogenous α-amino acid in place of the desired ncAA.^42–44^ Many factors can influence the misincorporation rate, including inefficient transport of the ncAA into cells, imperfect specificity of the aaRS enzyme for the ncAA over natural α-amino acids, differences in EF-Tu-mediated delivery of the mis-acylated tRNA to the ribosome, and slower rates of accommodation, elongation, or translocation of mis-acylated tRNAs.^43,64,65^ Misincorporation at an amber stop codon can also arise from near-cognate suppression, in which natural (and correctly acylated) tRNAs pair imperfectly with a UAG codon and direct the incorporation of one or more natural α-amino acids in place of the ncAA. The frequency of near-cognate suppression events depends on many factors, including the concentration of charged suppressor tRNA relative to near-cognate tRNAs and the presence of RF1, with higher occurrence of near-cognate suppression when the level of charged tRNA is low and RF1 is absent.^42^ It has been found that G/U mismatch as well as the 3^rd^-base wobble mismatch are the most common errors during codon recognition, followed by some other single-base mismatches (A/C, U/U and U/C).^66,67^ It follows that the aminoacyl-tRNA of tyrosine (both codons), glutamine (CAG codon), tryptophan, lysine (AAG codon), and glutamic acid (GAG codon) are likely near-cognate suppressors of UAG, in agreement with observations here and elsewhere.^68^ Additionally, phenylalanine and tyrosine misincorporations are also expected due to tRNA mischarging for their structural similarities to the ncAAs (pAzF, pBrF, and pCNF).

We used high-resolution LC-MS/MS to evaluate and compare the strain-dependent fidelity of ncAA incorporation at position 2 of sfGFP using pCNFRS (**Figure 3B** and **Supplementary Figure 6**). In each case, sfGFP samples were isolated from each strain after 4 h using IMAC, denatured, reduced and alkylated with iodoacetamide, and finally digested with Glu-C. The peptide fragments so-generated were analyzed on an LC-MS/MS system composed of an Agilent 1290-II HPLC and a Thermo Fisher Q Exactive Biopharma mass spectrometer. The mass spectrometry data were searched against the sfGFP sequence using MassAnalyzer (an in-house developed program, available in Biopharma Finder from Thermo Fisher) for potential amino acid substitutions.^66^

The LC-MS/MS data revealed that the yield of sfGFP containing the desired ncAA at position 2 was both strain and ncAA dependent (**Supplementary Figure 6**). As expected from the reported substrate preferences of pCNFRS, the highest and lowest fidelity were observed with pBrF and pCNF, respectively.^21^ The yield of sfGFP containing pBrF at position 2 ranged from a high of 99.7% (in Top10 cells) to a low of 99.4% (BL21 cells). With pAzF, the range was slightly greater, from a high of 97.6% (Vmax X2) to a low of 91.7% (C321.ΔA.opt). The lowest fidelity was observed for the incorporation of pCNF. Here, the fidelity was low (between 54.5% and 61.6%) in all strains except Vmax X2, where the fidelity was 91.8%. It is notable that the most significant impurities, regardless of strain, contained Phe in place of the desired ncAA, although in certain cases a significant amount of Tyr was also detected (**Supplementary Figure 6)**. The improved fidelity of sfGFP containing a ncAA at position 2 when produced in Vmax X2 could be due to many factors, including differences in ncAA permeability, aaRS activity, tolerance of Vmax X2 EF-Tu to mis-acylated tRNAs, even the relative activity of Vmax X2 RF1. Nevertheless, if the goal is a homogeneous and uniquely modified ncAA-containing protein in high yield and purity, Vmax X2 outperformed other strains, even those that funnel GCE through a single UAG channel. This statement is especially true in the case of ncAA possessing moderate to low specific activity, which could obviate the need for directed evolution to improve specific activity further.

We note that in the previous experiments, the plasmids used to express the aaRS/tRNA pair and sfGFP in Vmax X2 cells were identical to the plasmids used in BL21, but those used in C321 and Top10 cells were different. In the case of Vmax X2 and BL21, the aaRS/tRNA pair was under the control of an arabinose-dependent promoter and sfGFP expression was under control of the T7 promoter. However, as C321 and Top10 cells lack T7 RNA polymerase, in this case, aaRS/tRNA expression was under control of a tac promoter and sfGFP was under control of an arabinose promoter. We wondered about the extent to which these differences in promoter/inducer identity affected the relative yield of sfGFP in each cell line (**Supplementary Figure 7**). To evaluate this question, we made use of a plasmid in which the expression of sfGFP and the aaRS/tRNA pair expression were under control of T5 promoter and arabinose promoters, respectively - both of which are compatible with all cell types examined here.^45^ In this case, the plasmid encoded sfGFP carried a UAG codon at position 151. When under the control of a T5 promoter and in the presence of pBrF, although the OD_600_ of Vmax X2 cells was highest, the value of 528 nm emission at the 4 h timepoint was highest for BL21 and C321.ΔA.exp cells, lowest for C321.ΔA.opt and Top10 cells, and intermediate for Vmax X2 cells (**Supplementary Figure 7A**). The isolated yield of 151TAG sfGFP was comparable whether expression was performed in BL21, C321.ΔA.exp, or Vmax X2 – approximately 100 mg/L of culture. These yields were at least 2-fold lower than those obtained in Vmax X2 cells when sfGFP expression was under control of a T7 promoter (**Supplementary Figure 7B and 7C)**.

For many ncAA applications relating to fundamental research or the preparation of homogeneous antibody-drug conjugates (ADCs), one ncAA per polypeptide chain is generally sufficient. For others, such as the design of sequence-defined protein materials, multiple copies of one (or more) ncAA may be desired.^69^ These applications push the limits of genetic code expansion, as the isolated yields of such materials from standard *E. coli* strains can be low due to the increased frequency of RF1-mediated termination events.^45^ Strains that lack RF1, such as B95 and C321 derivatives, have been reported to support greatly improved yields of model proteins containing multiple copies of a single ncAA such as pAzF.^39,41,45^ Likewise, cell-free expression systems derived from these strains have been utilized.^55,70,71^ These systems are able to produce appreciable amounts of material containing up to 40 non-canonical amino acids,^70^ but fidelity is evaluated only rarely.

We wondered whether Vmax X2 would also provide advantages in yield or purity for expression of proteins whose coding sequences contained multiple UAG codons. Thus, we examined the strain-dependent yield and purity of sfGFP containing pBrF at five positions with sfGFP (S2, D36, K101, E132, and D190). All of these positions have been shown previously to accept one of more ncAAs.^71,72^ As before, all strains were grown under their own optimized conditions and in the presence of 0.5 mM pBrF; both OD_600_ and fluorescence at 528 nm were monitored as a function of time (**Figure 4A and B**). As observed previously, Vmax X2 cells grew fastest (**Figure 4A**) and there was no direct correlation between the 528 nm emission value at 4 h and the isolated yield. Although the average 528 nm emission of C321.ΔA.exp cells was higher than that of the Vmax X2 growth (**Figure 4D)**, the isolated yield of sfGFP from Vmax X2 was significantly higher (**Figure 4E**). The fidelity of sfGFP containing five copies of pBrF was also higher in Vmax X2 (>98.7%) than in C321.ΔA.exp (**Supplementary Figure 7**). Other detected amino acids include phenylalanine, glutamine (C321 strain only), tyrosine, lysine, leucine/isoleucine, and proline (**Supplementary Figure 8**). None of the other E. coli strains generated sufficient material for detailed LC-MS/MS analysis.

**Legend for Figure 4.**
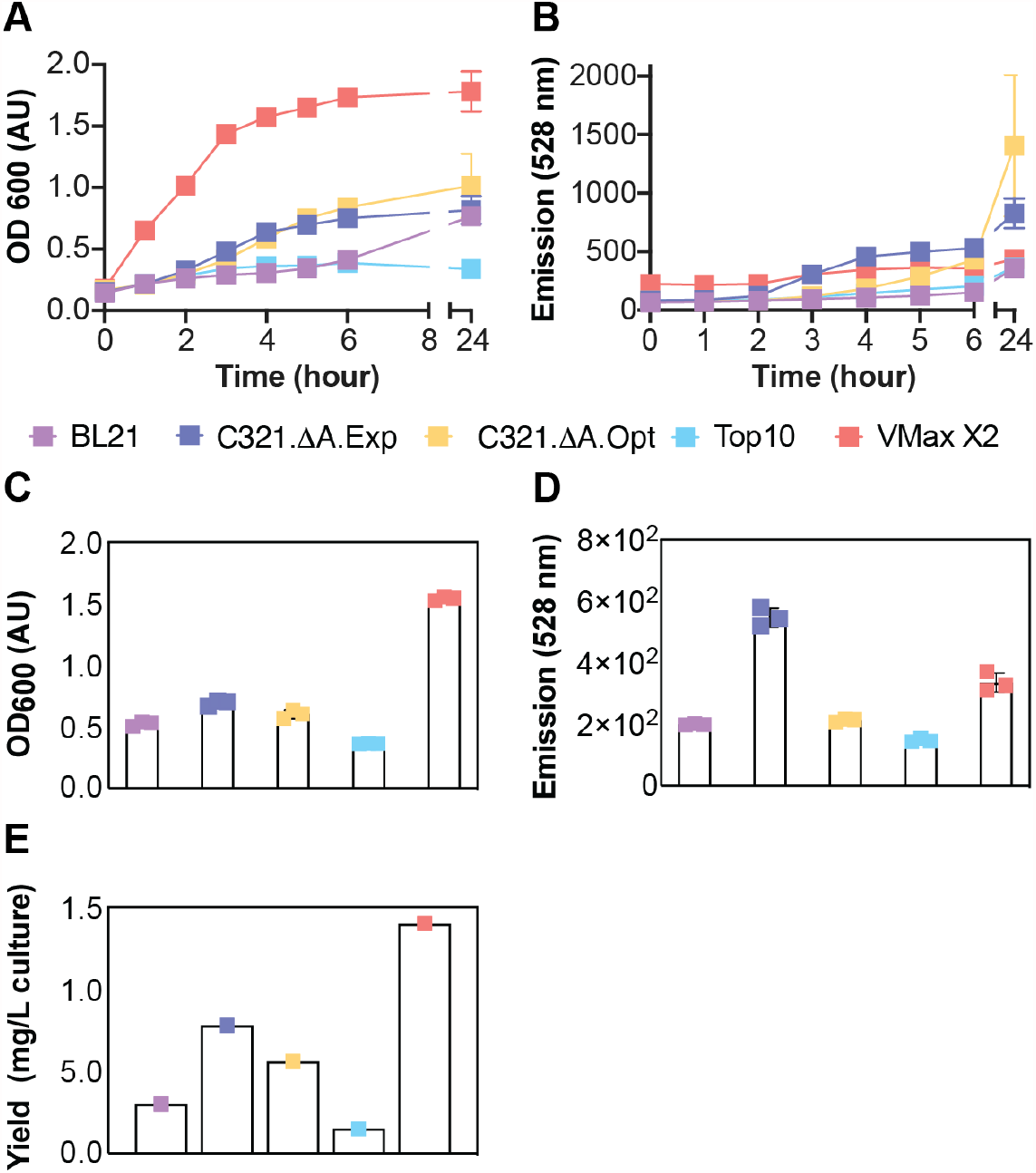
Yield of sfGFP containing five copies of pBrF ncAA when expressed in Vmax X2 versus *E. coli* strains. (A) Time-dependent growth curves; (B) Time-dependent increase in 528 nm fluorescence; (C) OD_600_; (D) 528 nm fluorescence; and (E) isolated yields after 4 h. Vmax X2 and BL21 cells were transformed with pET-5XTAG-sfGFP and pEVOL-CNF, whereas Top10 and C321 cells were transformed with pET-5XTAG-sfGFP and pULTRA-CNF (Top10, C321), to induce expression of sfGFP bearing a ncAA at five positions with sfGFP (S2, D36, K101, E132, and D190). After induction, cells were grown for 4 hours at 37°C (Vmax X2, BL21, Top10, C321.ΔA.exp) or 34°C (C321.ΔA.opt) in the presence of 0.5 mM pBrF.

In summary, here we show that Vmax X2 is capable of producing exceptional yields of soluble protein containing up to 5 ncAAs. The yields realized with Vmax X2 are up to 25-fold higher than those achieved using commercial expression strains (Top10 and BL21) and more than 10-fold higher than those achieved using two different genomically recoded *E. coli* strains that lack endogenous UAG stop codons and have been optimized for improved fitness and preferred growth temperature (C321.ΔA.opt and C321.ΔA.exp, Addgene strains #87359 and #49018).^39,40^ In addition to high yields, Vmax X2 cells also generate proteins with significantly lower levels of mis-incorporated natural α-amino acids at the UAG-programmed position, especially in cases in which the ncAA is only a moderate substrate for the chosen aaRS. Thus, use of Vmax X2 can obviate the need for time-consuming directed evolution experiments to improve specific activity of highly desirable substrates.

## Acknowledgements

This work was supported by the Center for Genetically Encoded Materials (C-GEM), an NSF Center for Chemical Innovation (NSF <u>2002182</u>). O.A. was supported in part by Agilent Technologies as an Agilent Fellow. S.S. was supported by an NSF Predoctoral Fellowship (NSF 1752814) and an NIH Chemistry-Biology Interface Training Grant (T32GM067543).

## I. Supplementary Figures

**Supplementary Figure 1.**
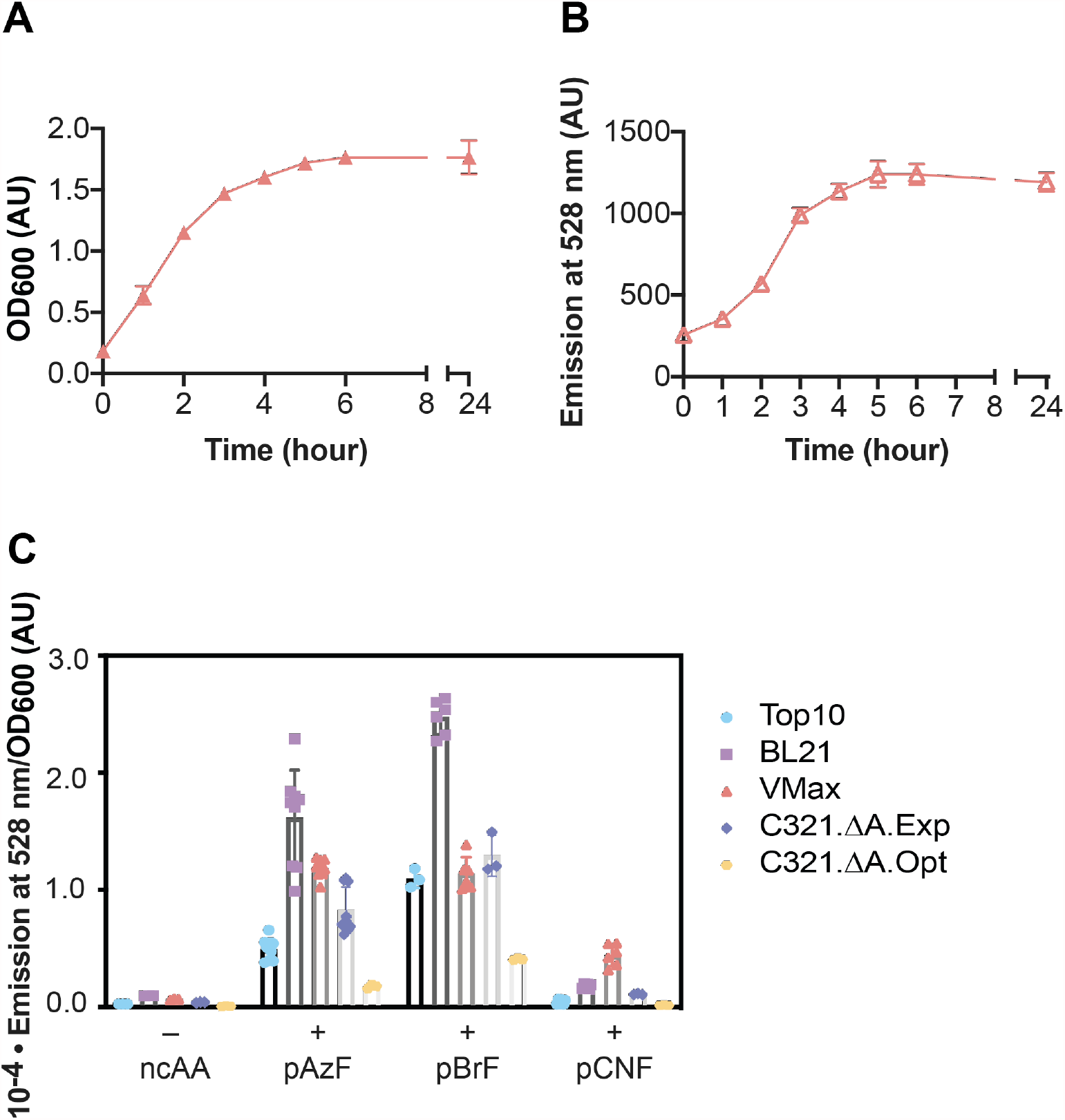
Growth and sfGFP expression of Vmax X2 cells transformed with pET-S2TAGsfGFP and pEVOL-mmPyl. Vmax X2 cells were transformed with pET-S2TAGsfGFP^1^ and pEVOL-mmPyl^2^ to induce expression of sfGFP bearing a ncAA at the second position of sfGFP. After induction, cells were incubated at 37°C in Brain-Heart Infusion broth supplemented with V2 salts for 24 hours in the presence of 10 mM BocK. Time-dependent changes in (A) OD_600_ and (B) fluorescence emission at 528 nm. (C) Plots comparing the OD-normalized 528 nm fluorescence of each strain at the 4 h time point in the presence or absence of pAzF, *p*-bromo-L-phenylalanine (pBrF), or *p*-cyano-L-phenylalanine (pCNF).

**Supplementary Figure 2.**
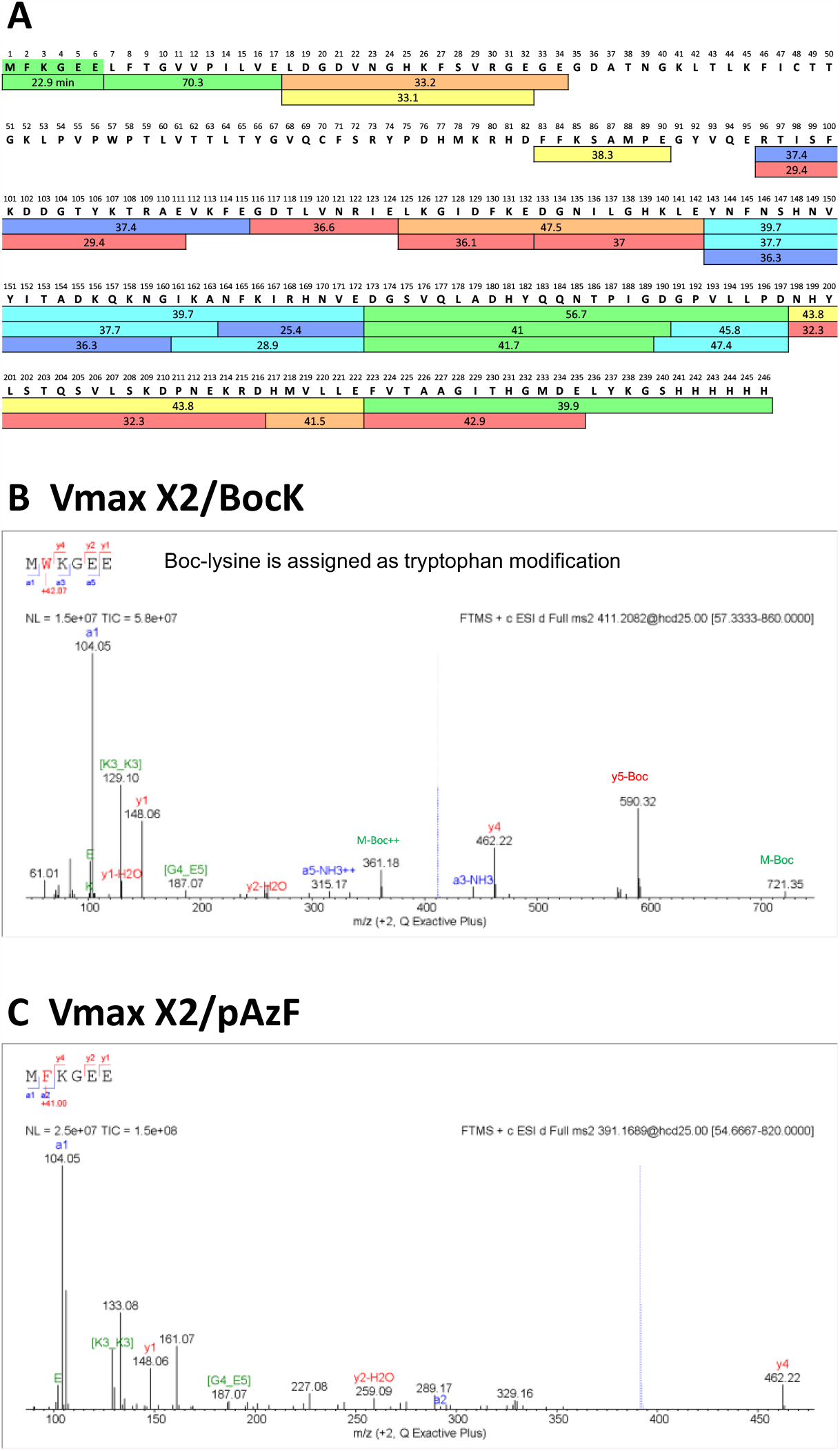
LC-MS/MS analysis of sfGFP expressed in Vmax X2. Vmax X2 cells were transformed with pET-S2TAGsfGFP^1^ and either pEVOL-CNF^3^ or pEVOL-mmPyl^2^ to induce expression of sfGFP bearing a ncAA at position two. Cells were induced and incubated for 4 hours at 37°C in the presence of 0.5 mM pAzF (pCNFRS) or 10 mM BocK (MmPylRS), purified using IMAC, and submitted for LC-MS/MS analysis. (A) Sequence of wild type sfGFP illustrating the peptide fragments obtained after digestion with Glu-C and their retention times. Colors from red to blue represent decreasing signal intensity. (B) MS/MS identification of the major N-terminal peptide derived from sfGFP isolated from Vmax X2 cells transformed with pET-S2TAGsfGFP and pEVOL-mmPyl and incubated for 4 h in the presence of 10 mM BocK. (C) MS/MS identification of the major N-terminal peptide derived from sfGFP isolated from Vmax X2 cells transformed with pET-S2TAGsfGFP and pEVOL-CNF and incubated for 4 h in the presence of 0.5 mM 4-azido-L-phenylalanine (pAzF). The fidelity of BocK incorporation at position 2 in Vmax X2 cells was ∼99%. Other amino acids that could be detected at position 2 include W (0.46%), L/I (0.31%), and Y (0.06%).

**Supplementary Figure 3.**
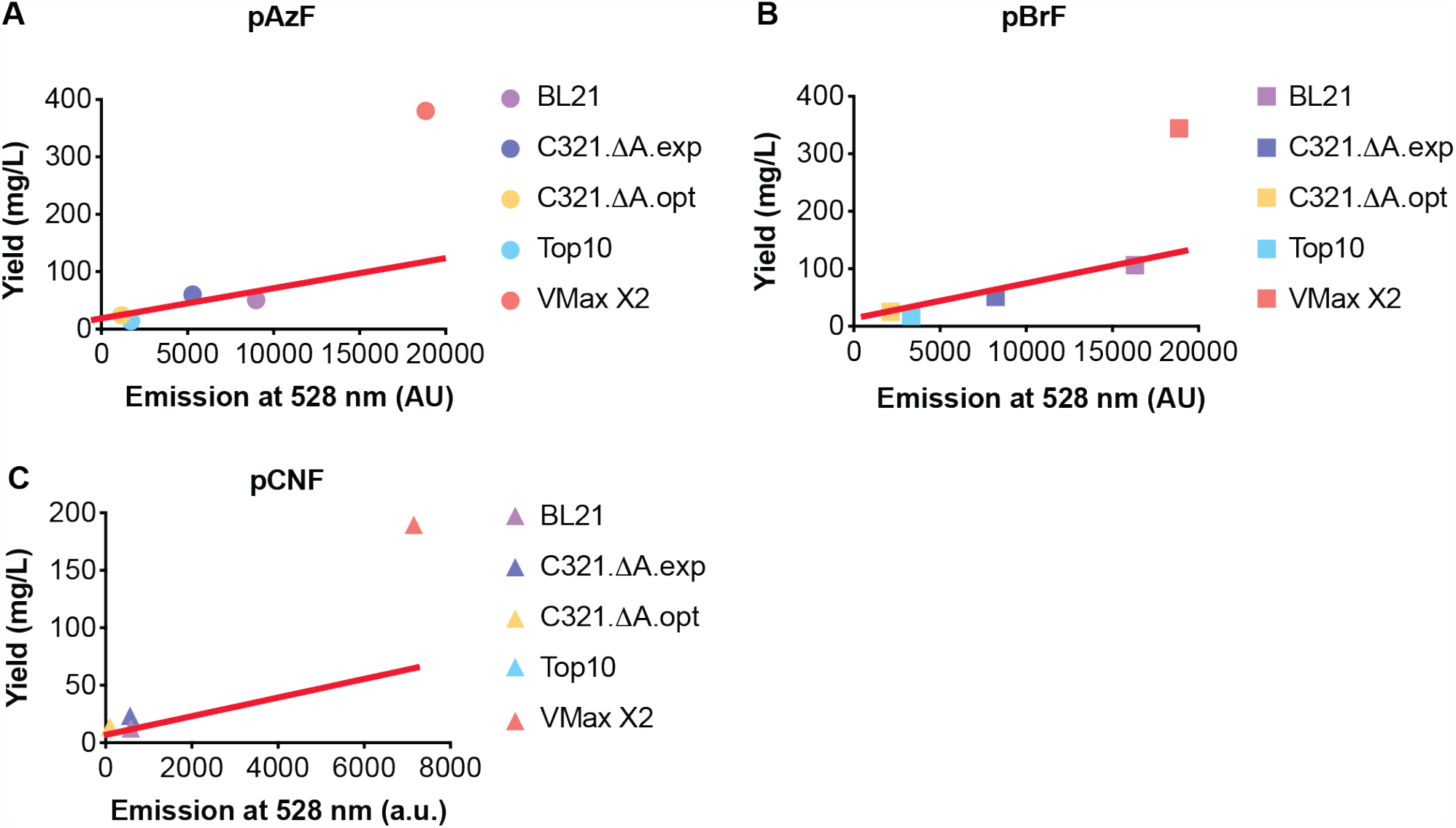
Isolated yield of sfGFP containing a ncAA at position 2 is higher than expected based on unit 528 nm emission. Plot of the isolated yield of sfGFP containing (A) pAzF; (B) pBrF; or (C) pCNF at position 2 versus the absorbance of the cell growth 4 h after induction. The red line shows the yield expected if yield correlated directly with unit 528 nm emission at 4 h.

**Supplementary Figure 4.**
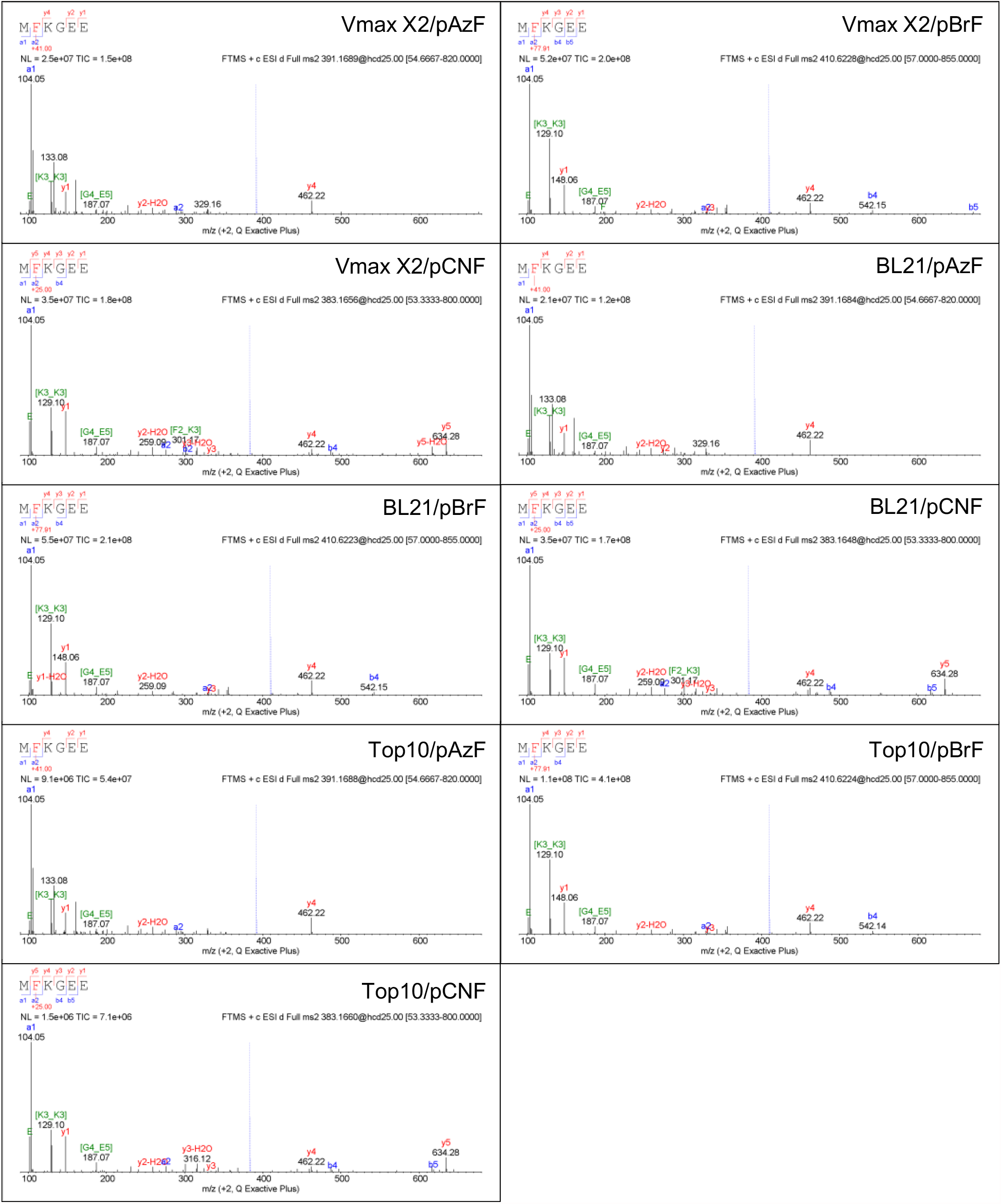
MS/MS identification of the N-terminal sfGFP peptide MXKGEE containing pAzF, pBrF, or pCNF at position 2 when produced in Vmax X2, BL21, or Top10 cells. The N-terminal peptide MXKGEE was generated by Glu-C digestions of sfGFP samples obtained from the indicated strain in growths containing the indicated ncAA.

**Supplementary Figure 5.**
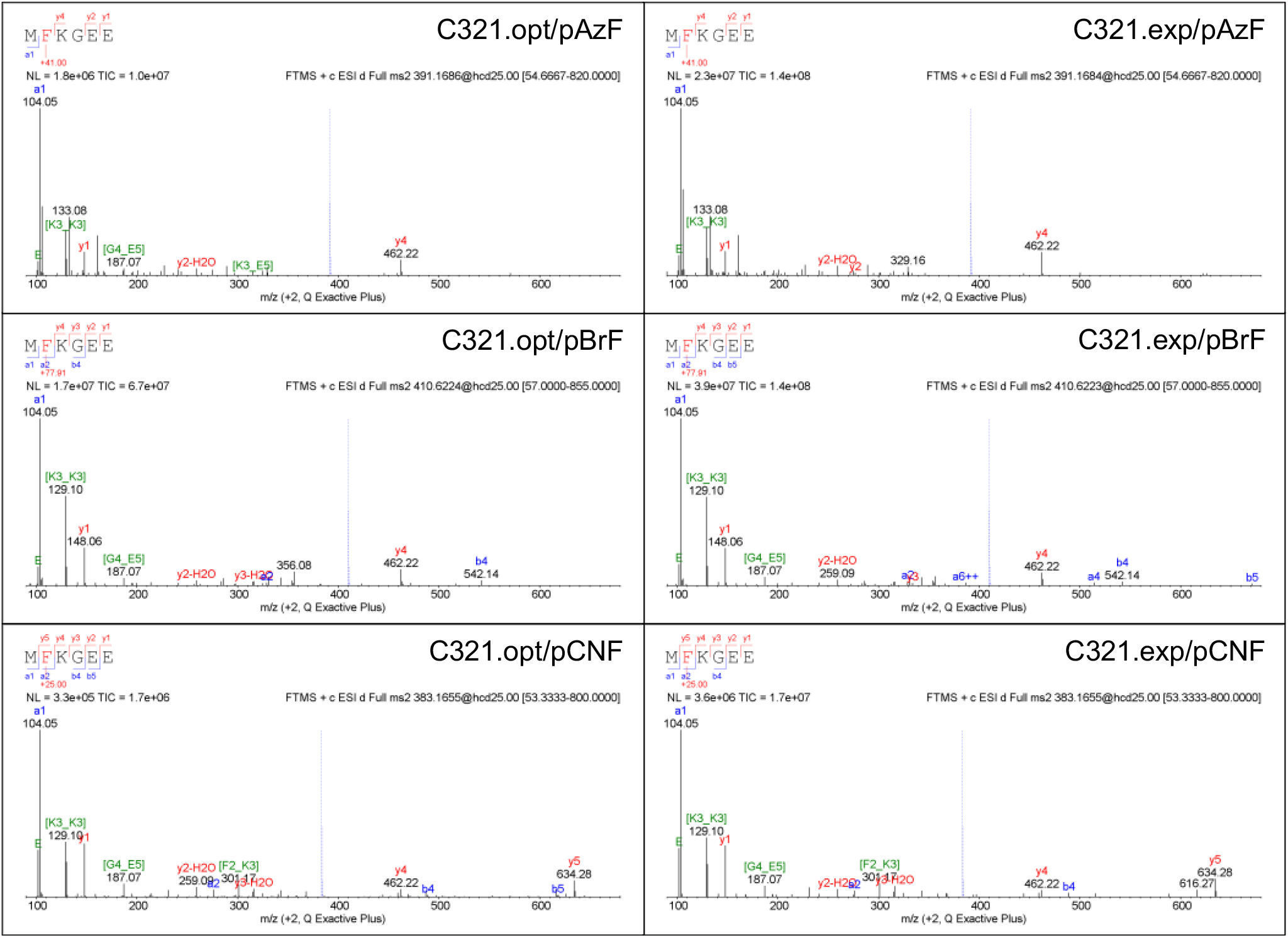
MS/MS identification of the N-terminal sfGFP peptide MXKGEE containing pAzF, pBrF, or pCNF at position 2 when produced in C321.ΔA.exp and C321.ΔA.opt cells. The N-terminal peptide MXKGEE was generated by Glu-C digestions of sfGFP samples obtained from the indicated strain in growths containing the indicated ncAA.

**Supplementary Figure 6.**
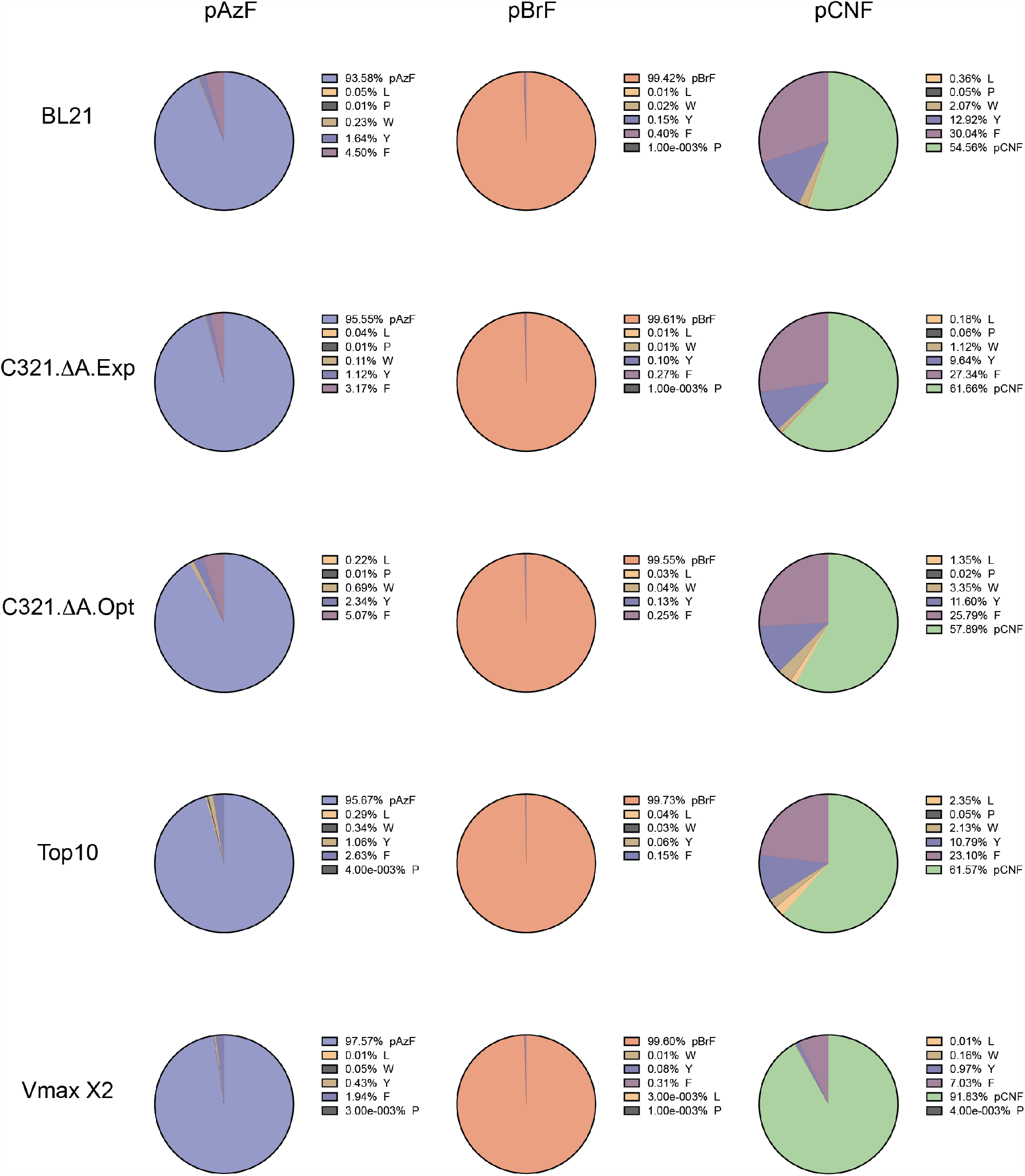
Pie charts plotting the distribution of amino acids incorporated at position 2 of sfGFP when expressed in the indicated strain. His-tagged proteins were isolated from growths of the indicated cells and digested with Glu-C. The relative abundance of the N-terminal sfGFP peptide fragments comprising the sequence MXKGEE were analyzed via LC-MS/MS to determine the identity of the amino acid at position 2. After sequence identification, relative amounts were calculated by integrating the area under the peak for each extracted ion chromatogram.

**Supplementary Figure 7.**
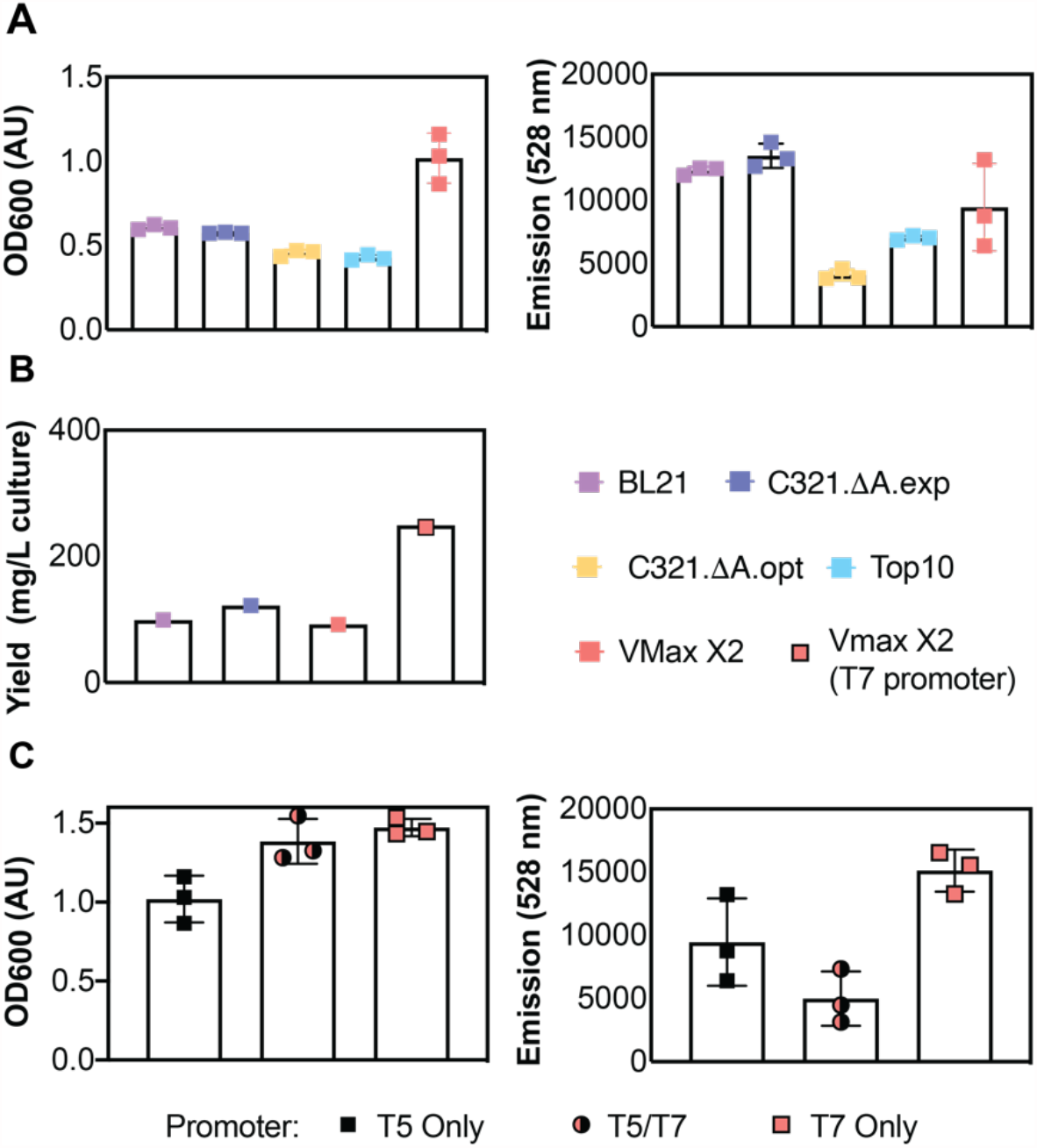
Comparison of GCE in Vmax X2 *versus* traditional (Top10, BL21) and genomically recoded (C321)^41,42^ using plasmids under the control of T5 and/or T7 promoters. (A) Plot of the OD_600_ and emission at 528 nm of each cell growth at the 4 h timepoint. All cells were transformed with pET-22B-151TAG sfGFP and pEVOL-CNF to induce expression of sfGFP bearing a ncAA at position 151 of sfGFP. After induction, cells were grown for 4 hours at 37°C (Vmax X2, BL21, Top10, C321.ΔA.exp) or 34°C (C321.ΔA.opt) in the presence of 0.5 mM pBrF. (B) Plot of the isolated yield of sfGFP obtained from each growth after 4 h incubation. The isolated yield of 2TAG sfGFP when expression is under control of the T7 promoter is shown for comparison. (C) Plots comparing the OD_600_ and 528 nm fluorescence Vmax X2 cells grown 0.5 mM pBrF as a function of promoter identity.

**Supplementary Figure 8.**
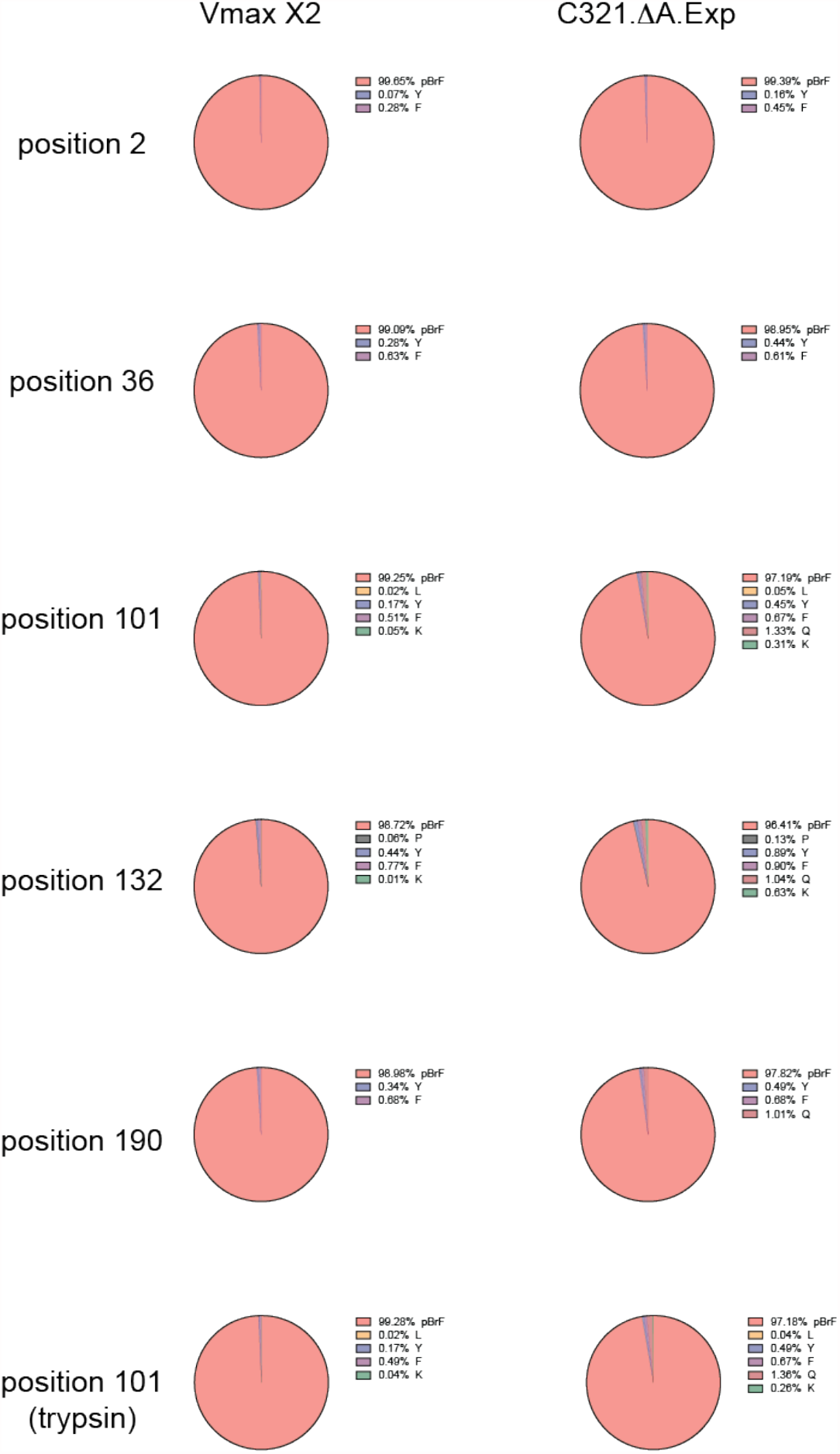
Pie charts plotting the distribution of amino acids incorporated at positions 2, 36, 101, 132, and 190 of sfGFP expressed in Vmax X2 or C321.ΔA.exp cells. His-tagged proteins were isolated from growths of the indicated cells and digested with Glu-C and/or trypsin.

## II. Supplementary Tables

**Supplementary Table 1.** Isolated yields (mg/L) of ncAA-containing sfGFP isolated from Vmax X2, BL21, Top10, C321.ΔA.exp, and C321.ΔA.opt cells after 4 h incubation. VMax X2 and BL21 cells expressed sfGFP and pCNFRS under the control of T7 and arabinose promoters, respectively. Top10, C321.ΔA.exp, and C321.ΔA.opt cells expressed sfGFP and pCNFRS under the control of arabinose and tac promoters, respectively.

| strain/<br>ncAA | VMax X2<br>(ratio) | BL21 | C321.ΔA.exp | C321.ΔA.opt | Top10 | Promoter<br>(sfGFP/aaRS) |
| --- | --- | --- | --- | --- | --- | --- |
| pAzF | 355<br>(5-fold to<br>25-fold) | 66 |  |  |  | T7/Arabinose |
|  |  | 42 | 62 | 18 | 14 | Arabinose/Tac |
| pBrF | 226<br>(2-fold to<br>12-fold) | 107 |  |  |  | T7/Arabinose |
|  |  |  | 25 | 18 | 18 | Arabinose/Tac |
|  | 92 | 99 | 122 |  |  | T5/Arabinose |
| pCNF | 115<br>(8-fold to<br>13-fold) | 9 |  |  |  | T7/Arabinose |
|  |  |  | 19 | 14 | 9 | Arabinose/Tac |

**Supplementary Table 2.** Plasmids used in this study.

| Plasmid Name | Resistance | Ori | ORF 1 |  | ORF 2 |  |
| --- | --- | --- | --- | --- | --- | --- |
|  |  |  | Promoter | Gene | Promoter | Gene |
| pET-S2TAG-sfGFP | Carb | pBR322 | T7 | S2TAG-sfGFP |  |  |
| pBAD-S2TAG-sfGFP | Carb | pBR322 | P<sub>BAD</sub> | S2TAG-sfGFP |  |  |
| pET22b-T5-151TAG-sfGFP | Carb | pBR322 | T5 | 151TAG-sfGFP |  |  |
| pET22b-T5/7-151TAG-sfGFP | Carb | pBR322 | T5/T7 | 151TAG |  |  |
| pET22b-T7-151TAG-sfGFP | Carb | pBR322 | T7 | 151TAG |  |  |
| pET-5xTAG-sfGFP | Carb | pBR322 | T7 | 5xTAG-sfGFP |  |  |
| pBAD-5xTAG-sfGFP | Carb | pBR322 | P<sub>BAD</sub> | sfGFP |  |  |
| pULTRA-CNF | Spec | CloDF13 | tac | pCNFRS |  |  |
| pEVOL-CNF | Cm | p15A | P<sub>BAD</sub> | pCNFRS | glnS | pCNFRS |

## III. Supplementary Methods

### Bacterial Strains

Vmax X2, BL21 (DE3), and Top10 cells were purchased from Codex DNA, NEB (Catalog # C2527), and ThermoFisher (Catalog # C404010) respectively. C321.∆A.Opt and C321.∆A.Exp were gifts from George Church (Addgene plasmids #87359 and #49018).

### Amino Acids

pAzF, pBrF, and pCNF were purchased from Chem-Impex International (Catalog # 06162, 04086, and 04110). BocK was purchased from Sigma-Aldrich (SKU 349661).

### General Methods

The following antibiotic concentrations were used: carbenicillin, 100 μg/mL (*E. coli)* or 12.5 μg/mL (VMax X2); chloramphenicol 25 μg/mL; spectinomycin 50 μg/mL. Additionally, C321.ΔA.opt and C321.ΔA.exp starter cultures were grown in the presence of 15 μg/mL gentamicin or 25 μg/mL Zeocin^™^ (ThermoFisher) respectively.

### Transformation protocols

Vmax X2 and *E. coli* (BL21, C321, Top10) cells were transformed in accordance with manufacturer protocols with some modifications as follows. Frozen stocks were thawed on ice. Upon thawing, 1 µL of plasmid (see below) encoding sfGFP and the orthogonal synthetase was added. After a 30 min incubation on ice, cells were heat shocked for 45 (Vmax X2) or 90 s (BL21, C321, Top10) and put back on ice for 2 minutes. 550 µL of Vmax recovery media (Vmax X2) or SOB media was added and cells were recovered at 34°C (C321.ΔA.opt) or 37°C for one (*E. coli*) or four hours (Vmax X2) before plating. For 2TAG-sfGFP expression, Vmax X2 and BL21 cells were transformed with pEVOL-CNF and pET-S2TAG-sfGFP, while C321 and Top10 cells were transformed with pULTRA-CNF and pBAD-S2TAG-sfGFP. pEVOL-mmPyl was used in place of pEVOL-CNF for expression of sfGFP containing BocK at the 2^nd^ position. For 5XTAG-sfGFP expression Vmax X2 and BL21 cells were transformed with pEVOL-CNF and pET-5XTAG-sfGFP, C321 and Top10 cells were transformed with pULTRA-CNF and pBAD-5XTAG-sfGFP. For 151TAG-sfGFP expression, all strains were transformed with pEVOL-CNF and pET22b-151TAG-sfGFP.

### Expression of sfGFP variants

Starter cultures were grown for 3 hours (Vmax X2) or overnight (*E. coli*) in 25 mL of BHI (Teknova, Catalog #B9505) + v2salts (204 mM NaCl, 23.2 mM MgCl_2_, 4.2 mM KCl) or 10 mL of LB Miller (AmericanBio, Catalog #AB01201) supplemented with antibiotics at 34°C (C321.A.opt) or 37°C. To maximize aeration growth rate, starter cultures for Vmax X2 cells were grown in a baffled flask. Following the initial incubation period, starter cultures were diluted 1:100 into 25 mL BHI + v2salts or LB supplemented with 0.5 mM (pAzF, pBrF, pCNF) or 10 mM (BocK) ncAA. Once cultures reached an OD of 0.5, protein expression was induced by addition of 1 mM IPTG and 0.2% arabinose.

### Purification of sfGFP variants

Cultures were centrifuged for 15 minutes at 10,000 x g and 4°C. Pellets were resuspended in 15 mL of lysis buffer (50 mM sodium phosphate (pH 8), 300 mM NaCl, 20 mM imidazole,) supplemented with 1 tablet cOmplete, mini EDTA-free protease inhibitor cocktail (Sigma-Aldrich, St. Louis, MO) and sonicated for 5 minutes total (30s on, 30s off) at 30% duty cycle. Following sonication, the soluble fraction was isolated by centrifugation of the lysate for 25 minutes at 10,000 x g and 4°C. The supernatant was isolated and incubated with 500 μL of Ni-NTA (Qiagen, Catalog # 30230) resin for an hour at 4°C. Slurry was poured onto a gravity flow column and the resin was washed with 15 mL of lysis buffer following drainage of the flowthrough. Bound protein was then eluted by the addition of 3.5 mL of elution buffer (50 mM sodium phosphate (pH 8), 250 mM imidazole). For quantification and MS analysis, the eluent was buffer exchanged into 50 mM sodium phosphate utilizing a PD-10 column.

### Intact Protein Mass Spectomery Analysis

LC/MS analysis was performed on an Agilent 1290 Infinity II HPLC connected to an Agilent 6530B QTOF AJS-ESI. 1 μg of protein was injected onto a Poroshell 300SB-C8 column (2.1 x 75 mm, 5-Micron, room temperature, Agilent) using a linear gradient from 5 to 75% acetonitrile over 9.5 minutes with 0.1% formic acid as the aqueous phase after an initial hold at 5% acetonitrile for 0.5 min (0.4 mL/min). The following parameters were used during acquisition: Fragmentor voltage 175 V, gas temperature 300°C, gas flow 12 L/min, sheath gas temperature 350°C, sheath gas flow 12 L/min, nebulizer pressure 35 psi, skimmer voltage 65 V, Vcap 5000 V, 3 spectra/s. Intact protein masses were obtained via deconvolution using the Maximum Entropy algorithm in Mass Hunter Bioconfirm (V10, Agilent).

### Monitoring of cell growth and sfGFP expression over time

25 mL cultures were grown in 250 mL baffled flask and expression sfGFP variants and pCNFRS was induced as described previously. At each timepoint, 100 uL aliquots of each culture were transferred onto a black, clear bottom, 96-well plate. The OD_600_ and emission at 528 nm (ƛ_ex_= 485 nm) was measured in BioTek Synergy H1 microplate reader.

### Fidelity of ncAA incorporation by LC-MS/MS

To determine the fidelity of amino acid incorporation at position 2 of sfGFP, isolated sfGFP (13 to 72 µg, most at ∼25 µg) was denatured with 6 M guanidine in a 0.15 M Tris buffer at pH 7.5, followed by disulfide reduction with 8 mM dithiothreitol (DTT) at 37 °C for 30 min. The reduced sfGFP was alkylated in the presence of 14 mM iodoacetamide at 25 °C for 25 min, and then quenched using 6 mM DTT. The reduced/alkylated protein was exchanged into ∼50 µL of 0.1 M Tris buffer at pH 7.5 using a Microcon 10-KDa membrane, followed by addition of 2.5 to 7.0 µg endoproteinase Glu-C (in a 0.5 µg/µL solution) directly to the membrane to achieve an enzyme-to-substrate ratio of at least 1:10. After 3 hours at 37°C, the digestion was quenched with an equal volume of 0.25 M acetate buffer (pH 4.8) containing 6 M guanidine. Peptide fragments were collected by spinning down through the membrane and subjected to LC-MS/MS analysis.

To determine the fidelity amino acid incorporation at the remaining 4 positions, isolated sfGFP was also digested by trypsin, with the same procedure as described above, except trypsin was used in place of Glu-C, and digested was allowed to proceed for 1 hour instead of 3 hours.

LC-MS/MS analysis was performed on an Agilent 1290-II HPLC directly connected to a Thermo Fisher Q Exactive high-resolution mass spectrometer. Peptides were separated on a Waters HSS T3 reversed-phase column (2.1 × 150 mm) at 50°C with a 70-min acetonitrile gradient (0.5% to 35%) containing 0.1% formic acid in the mobile phase, and a total flow rate of 0.25 mL/min. The MS data were collected at 70,000 resolution, followed by data-dependent higher-energy collision dissociation (HCD) MS/MS at a normalized collision energy of 25%. Proteolytic peptides were identified and quantified on MassAnalyzer, an in-house developed program (available in Biopharma Finder from Thermo Fisher). The program performs feature extraction, peptide identification, retention time alignment, and relative quantitation in an automated fashion.

### Sequences

* Denotes a stop codon

#### 2TAG-sfGFP

M* KGEELFTGVVPILVELDGDVNGHKFSVRGEGEGDATNGKLTLKFICTTGKLPVPWPTLVTTLTYGVQCFSRYPDHMKRHDFFKSAMPEGYVQERTISFKDDGTYKTRAEVKFEGDTLVNRIELKGIDFKEDGNILGHKLEYNFNSHNVYITADKQKNGIKANFKIRHNVEDGSVQLADHYQQNTPIGDGPVLLPDNHYLSTQSVLSKDPNEKRDHMVLLEFVTAAGITHGMDELYKGSHHHHHH

#### 5XTAG-sfGFP

M* KGEELFTGVVPILVELDGDVNGHKFSVRGEGEG* ATNGKLTLKFICTTGKLPVPWPTLVTTLT YGVQCFSRYPDHMKRHDFFKSAMPEGYVQERTISF* DDGTYKTRAEVKFEGDTLVNRIELKGID FK* DGNILGHKLEYNFNSHNVYITADKQKNGIKANFKIRHNVEDGSVQLADHYQQNTPIG* GPVL LPDNHYLSTQSVLSKDPNEKRDHMVLLEFVTAAGITHGMDELYKGSHHHHHH

#### 151TAG-sfGFP

MSKGEELFTGVVPILVELDGDVNGHKFSVRGEGEGDATNGKLTLKFICTTGKLPVPWPTLVTTLTYGVQCFSRYPDHMKRHDFFKSAMPEGYVQERTISFKDDGTYKTRAEVKFEGDTLVNRIELKGIDFKEDGNILGHKLEYNFNSHNV* ITADKQKNGIKANFKIRHNVEDGSVQLADHYQQNTPIGDGPVLLPDNHYLSTQSVLSKDPNEKRDHMVLLEFVTAAGITHGMDELYKGSHHHHHH

#### pCNFRS

MDEFEMIKRNTSEIISEEELREVLKKDEKSALIGFEPSGKIHLGHYLQIKKMIDLQNAGFDIIIVLADLHAYLNQKGELDEIRKIGDYNKKVFEAMGLKAKYVYGSEWMLDKDYTLNVYRLALKTTLKRARRSMELIAREDENPKVAEVIYPIMQVNGAHYLGVDVAVGGMEQRKIHMLARELLPKKVVCIHNPVLTGLDGEGKMSSSKGNFIAVDDSPEEIRAKIKKAYCPAGVVEGNPIMEIAKYFLEYPLTIKRPEKFGGDLTVNSYEELESLFKNKELHPMDLKNAVAEELIKILEPIRKRL

#### mmPylRS

MDKKPLNTLISATGLWMSRTGTIHKIKHHEVSRSKIYIEMACGDHLVVNNSRSSRTARALRHHKYRKTCKRCRVSDEDLNKFLTKANEDQTSVKVKVVSAPTRTKKAMPKSVARAPKPLENTEAAQA QPSGSKFSPAIPVSTQESVSVPASVSTSISSISTGATASALVKGNTNPITSMSAPVQASAPALTKSQTDRLEVLLNPKDEISLNSGKPFRELESELLSRRKKDLQQIYAEERENYLGKLEREITRFFVDRGFLEIKSPILIPLEYIERMGIDNDTELSKQIFRVDKNFCLRPMLAPNLYNYLRKLDRALPDPIKIFEIGPCYRKESDGKEHLEEFTMLNFCQMGSGCTRENLESIITDFLNHLGIDFKIVGDSCMVYGDTLDVMHGDLELSSAVVGPIPLDREWGIDKPWIGAGFGLERLLKVKHDFKNIKRAARSESYYNGISTN L

